# Unbiased single-cell morphology with self-supervised vision transformers

**DOI:** 10.1101/2023.06.16.545359

**Authors:** Michael Doron, Théo Moutakanni, Zitong S. Chen, Nikita Moshkov, Mathilde Caron, Hugo Touvron, Piotr Bojanowski, Wolfgang M. Pernice, Juan C. Caicedo

## Abstract

Accurately quantifying cellular morphology at scale could substantially empower existing single-cell approaches. However, measuring cell morphology remains an active field of research, which has inspired multiple computer vision algorithms over the years. Here, we show that DINO, a vision-transformer based, self-supervised algorithm, has a remarkable ability for learning rich representations of cellular morphology without manual annotations or any other type of supervision. We evaluate DINO on a wide variety of tasks across three publicly available imaging datasets of diverse specifications and biological focus. We find that DINO encodes meaningful features of cellular morphology at multiple scales, from subcellular and single-cell resolution, to multi-cellular and aggregated experimental groups. Importantly, DINO successfully uncovers a hierarchy of biological and technical factors of variation in imaging datasets. The results show that DINO can support the study of unknown biological variation, including single-cell heterogeneity and relationships between samples, making it an excellent tool for image-based biological discovery.

## Introduction

The visual interpretation of cellular phenotypes through microscopy is widely used to drive biomedical discovery across a variety of questions in biology, including subcellular protein localization ^1, 2^, mitochondrial phenotypes ^3, 4^, cell cycle stages ^5, 6^, as well as chemical ^7^ and genetic perturbations ^8, 9^. Machine learning has the potential to unlock rich feature extraction of cellular morphological phenotypes that has thus far remained largely inaccessible to investigators ^10–12^, making single-cell morphological profiling among the current roster of single-cell omics in its own right ^13–16^. Indeed, deep learning now powers robust cell segmentation methods ^17–20^, as well as cell phenotyping using classification networks ^21, 22^.

Still, approaches to image analysis in microscopy remain specialized: many studies used to design bespoke workflows to measure one or two features ^23^, such as cell counts or cell size, disregarding much of the content present in modern microscopy. Image-based profiling workflows have been designed to quantify cellular morphology more broadly ^24, 25^, however, these are based on engineered features that may not capture all phenotypic variation encoded in images. Recently, supervised deep learning approaches have been successful in learning morphological features directly from data, yet this approach requires either extensive manual annotation efforts ^26, 27^ or some prior knowledge about the biology of interest ^28, 29^, yielding models that are highly application specific A general strategy for obtaining unbiased cellular phenotypes from images is still lacking. Such a strategy would make image-based phenotypes accessible to more studies, especially if there is no prior knowledge of phenotypic variation, opening the way to interrogate visual phenotypes in an unbiased way similar to how molecular phenotypes are investigated by gene expression.

Here, we show that DINO ^30^ (self-**di**stillation with **no** labels) is an effective self-supervised learning algorithm capable of obtaining unbiased representations of cellular morphology without the need for manual annotations or explicit supervision. DINO was originally designed for natural RGB images, and it has shown remarkable ability to discover meaningful factors of variation in image collections, and to achieve state-of-the-art performance in multiple vision problems ^31–34^. Furthermore, when DINO is used to train vision transformer (ViT) networks ^35^, the internal image representations exhibit emerging semantic properties that are not readily captured by convolutional networks, thanks to the transformers’ self-attention operations ^36, 37^.

These properties make DINO especially useful for cellular biology problems where the goal is to test hypotheses and uncover novel phenomena, and for powering various bioimage analysis applications ^31, 38, 39^. Our study demonstrates that DINO is a general, self-supervised approach to uncover meaningful biology from fluorescent microscopy images by analyzing three publicly available datasets with diverse technical specifications and biological goals (Figure 1). The results show that DINO features support a wide range of downstream tasks, from subcellular to population-level analyses, showing great potential for biological discovery.

**Figure 1.**
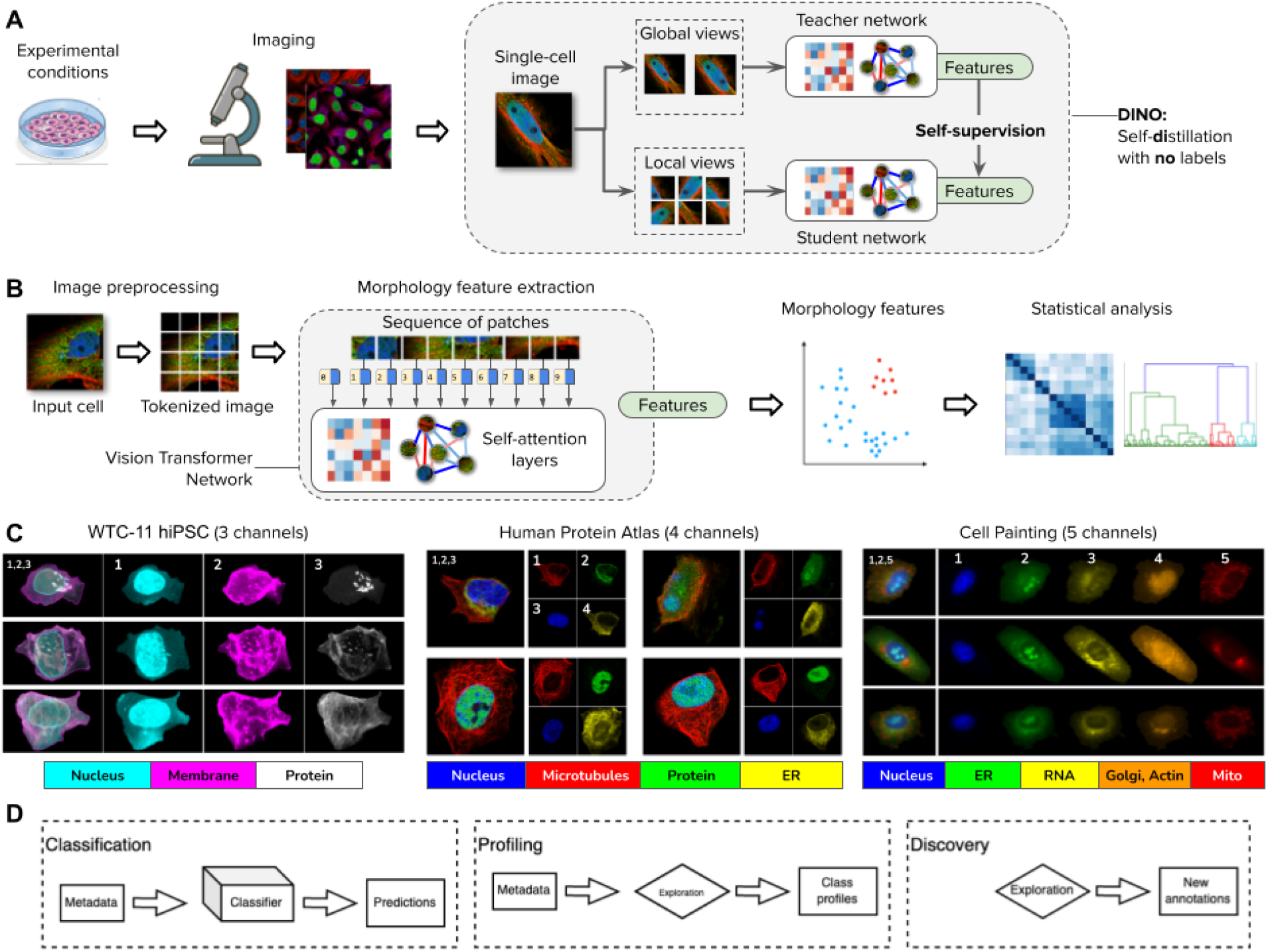
DINO: self-distillation with no labels, vision transformers, and datasets in this study. A) Illustration of the workflow for collecting images of experimental conditions and then analyzing cellular phenotypes with the DINO algorithm, which trains two neural networks: a teacher and a student network. DINO is trained on images only with a self-supervised objective that aims to match the features produced by the teacher (which observes random global views of an example image) with the features produced by the student (which observes random global and local views of the same example image) (Methods). B) Illustration of the processing pipeline using vision transformer networks trained with DINO. The image of a cell is first tokenized as a sequence of local patches, which the transformer processes using self-attention layers (Methods). The vision transformer network produces single-cell morphology feature embeddings, which are used for downstream analysis using other machine learning models or statistical tests. C) Example images of the datasets used in this study: the Allen Institute WTC-11 hiPSC dataset, the Human Protein Atlas, and a collection of Cell Painting datasets. D) Biological data analysis and downstream tasks enabled by DINO features in the datasets mentioned in C.

## Results

### Self-supervised vision transformers for single-cell images

The DINO ^30^ algorithm is a self-supervised learning strategy to transform images into data representations for quantitative analysis. This self-supervised strategy only requires images to create a meaningful feature space without using annotations or prior knowledge. DINO trains a feature extraction model using different views of the same image, which are shown independently to a student and a teacher network: during training their outputs should match (Figure 1A and Methods). By optimizing this simple rule, DINO learns features that capture semantic properties of images. In the case of images of single cells, these features are expected to encode biologically meaningful properties of cellular morphology.

The feature extraction model trained with DINO is a vision transformer (ViT), based on a neural network architecture with self-attention layers (Methods) originally developed for natural language processing ^35^ (Figure 1B). ViTs process an image by first decomposing it into a sequence of local patches and then treating them as if they were tokens in a sentence. In this way, ViTs capture long range dependencies between image regions by paying attention to all tokens simultaneously. This is in contrast to convolutional networks (CNNs), which are based on local feature analysis progressively expanded to cover larger spatial areas with pooling operations. While CNNs have been successful in many supervised image analysis problems, ViTs are more effective in self-supervised settings and scale efficiently to learn from large unannotated datasets. Self-supervised transformers also exhibit a unique ability to generate emerging properties and behaviors, which has been observed in images ^30, 40^ and natural language ^41^. Similarly, transformers have been exceptionally successful in genomics applications ^42, 43^ as well as protein folding prediction ^44^.

### DINO uncovers the biological structure of imaging experiments

We used DINO to analyze three publicly available imaging datasets (Figure 1C), and we observe that it discovers meaningful factors of variation directly from the images in all cases. The three datasets included in this study are the Allen Institute WTC-11 hiPSC (WTC11) dataset ^45^, the Human Protein Atlas (HPA) dataset ^1^, and a collection of Cell Painting datasets (CPC) ^29, 46, 47^. These datasets represent a diverse set of imaging techniques, cell lines, and experimental conditions, which have been used to study intracellular organization (WTC11), subcellular protein localization (HPA), and the effects of compound perturbations (CPC). All three datasets were pre-processed to obtain single-cell images using segmentation algorithms optimized for each case (Methods), and the DINO model was trained on single cells for each dataset separately to produce a morphology feature space.

This morphology feature space captures the phenotypic diversity of cells in the corresponding imaging study. Specifically, DINO identifies clusters of cellular structures in the WTC11 dataset (Figure 2A), clusters of cell lines in the HPA dataset (Figure 2D), and clusters of mechanisms of action (MOA) in the CPC dataset (Figure S1). These results illustrate how DINO encodes meaningful information without explicitly being trained to identify or detect such phenotypes.

**Figure 2.**
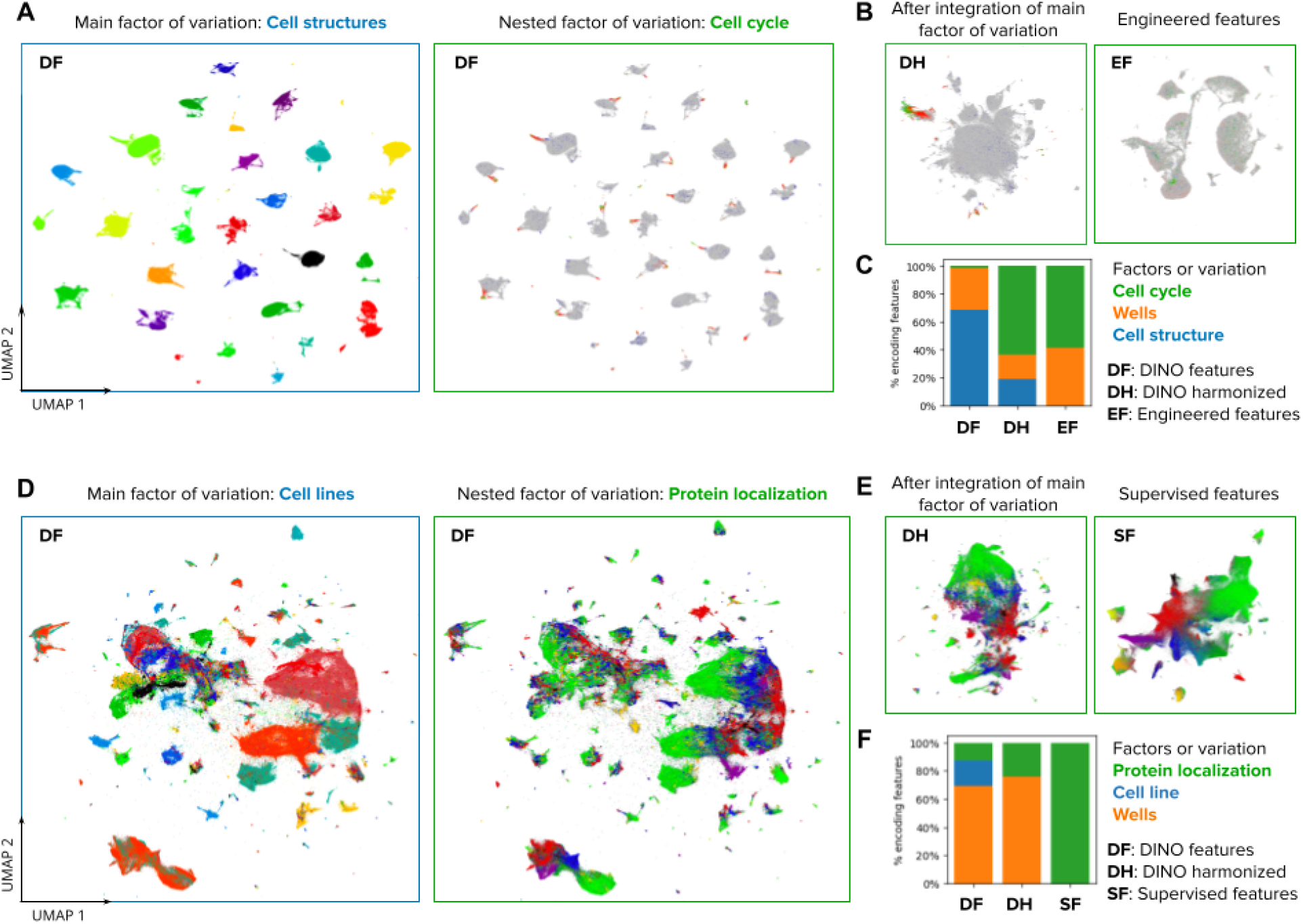
Visualizations of the morphology feature space automatically discovered by DINO. A) UMAP visualization of single-cell morphology features obtained with DINO for images in the Allen Institute WTC-11 hiPSC dataset. The two UMAP plots show the same data points, with the left plot coloring points by the endogenously tagged cell structure, and the right plot coloring points by the cell-cycle stage annotation. The title and frame colors match the legend in C for factors of variation. B) UMAP visualizations of the same single-cell images in A. The left-hand side plot shows DINO features integrated with Harmony over cell lines. The right-hand side plot shows engineered features used in the original study ^45^. C) Stacked-bar plot displaying the fraction of features that are strongly associated with three factors of variation annotated in the WTC-11 hiPCS dataset (colors). Three feature representation strategies are compared (bars). D) UMAP visualization of single-cell morphology features obtained with DINO for images in the Human Protein Atlas dataset. The left-hand side plot shows points colored by cell line, and the right-hand side plot shows points colored by protein localization labels. The title and frame colors match the legend in F for factors of variation. E) UMAP visualizations for the same single-cell images in D. The left-hand side plot shows DINO features integrated with Harmony over cell lines. The right-hand side plot shows features obtained with a supervised CNN from a top competitor in the weakly supervised single-cell classification challenge in Kaggle ^22^. F) Same as in C but with factors of variation annotated in the Human Protein Atlas dataset.

Importantly, we find that these morphology feature spaces also encode additional aspects of cellular variation in a hierarchically organized way: e.g., DINO locally encodes cell-cycle stages for each cellular structure cluster in WTC11 (Figure 2A), and protein localization variations for each cell line in HPA (Figure 2D).

The ability of DINO to encode diverse factors of variation in an unbiased way could enable various types of phenotypic analyses. Consistent with this, we transformed the DINO features using the single-cell data integration algorithm Harmony ^48^ to remove variation across the cellular structure clusters in WTC11 and the cell line clusters in HPA. Harmony integrates the corresponding clusters and preserves biological variation in both cases, resulting in feature spaces that respectively align cell-cycle stages across cell structures in the WTC11 dataset (Figure 2B), and that encode the continuum of protein localization variations in the HPA dataset (Figure 2E). Notably, the resulting integrated feature spaces qualitatively resemble the structure identified by specialized methods, using hand-engineered features (Figure 2B), or supervised methods trained specifically for protein localization classification (Fig. 2E).

To explore the hierarchy of the factors of variation encoded in DINO features quantitatively, we used a mutual-information based metric ^49^ to associate individual features with known data annotations (Methods). We find that DINO features capture a greater diversity of factors of variation compared to engineered features or supervised models, which specialize on specific variation according to the labels used for training (Figures 2C and 2F). Importantly, DINO captures not only biologically meaningful features, but also technical variation. Supervised learning regimes, in which feature extractors are often trained end-to-end, are prone to exploiting spurious correlations.^50^ However, similar to gene-expression data ^51^, DINO features can be transformed to correct for unwanted variation prior to downstream analyses (Figure 2B,E), potentially avoiding confounders. Nevertheless, effectively mitigating confounders remains an open challenge, which we do not investigate here.

### DINO features can predict expert-defined cellular phenotypes

A common task in bioimage analysis is the classification of phenotypes of interest according to expert-defined annotations. DINO features show highly competitive performance when used to train classifiers with manually annotated examples, even though the feature extractors are optimized independently with self-supervision and not trained to solve these tasks explicitly (Methods). We evaluated the performance of classifiers trained with DINO features on two classification tasks concerning manual annotation of single-cell phenotypes: cell-cycle stage classification in the WTC11 dataset, and protein localization in the HPA dataset. No fine-tuning of the DINO models was involved for training any of the classifiers.

For the cell-cycle stage classification task, we sampled a balanced subset of cells from WTC11 that represents six stages according to available annotations (Methods). We trained three Multi-Layer Perceptron (MLP) classifiers with the same architecture using the following features for comparison: engineered features available in the WTC11 dataset ^45^, DINO features from a model trained with natural images (ImageNet DINO ^52^), and DINO features from a model trained with WTC11 cells. The two sets of DINO features outperformed engineered features (Figure 3A and 3B), highlighting how self-supervision automatically captures information relevant for biological analyses. The DINO model trained on WTC11 achieves competitive results despite being trained with fewer images than ImageNet DINO (1.2 vs 0.2 million, 0.9% improvement over engineered features). ImageNet DINO seems to encode cell cycle variation more prominently (Figure S2) because the nucleus channel is processed separately, partially avoiding confounders and obtaining superior performance (18% relative improvement over engineered features).

**Figure 3.**
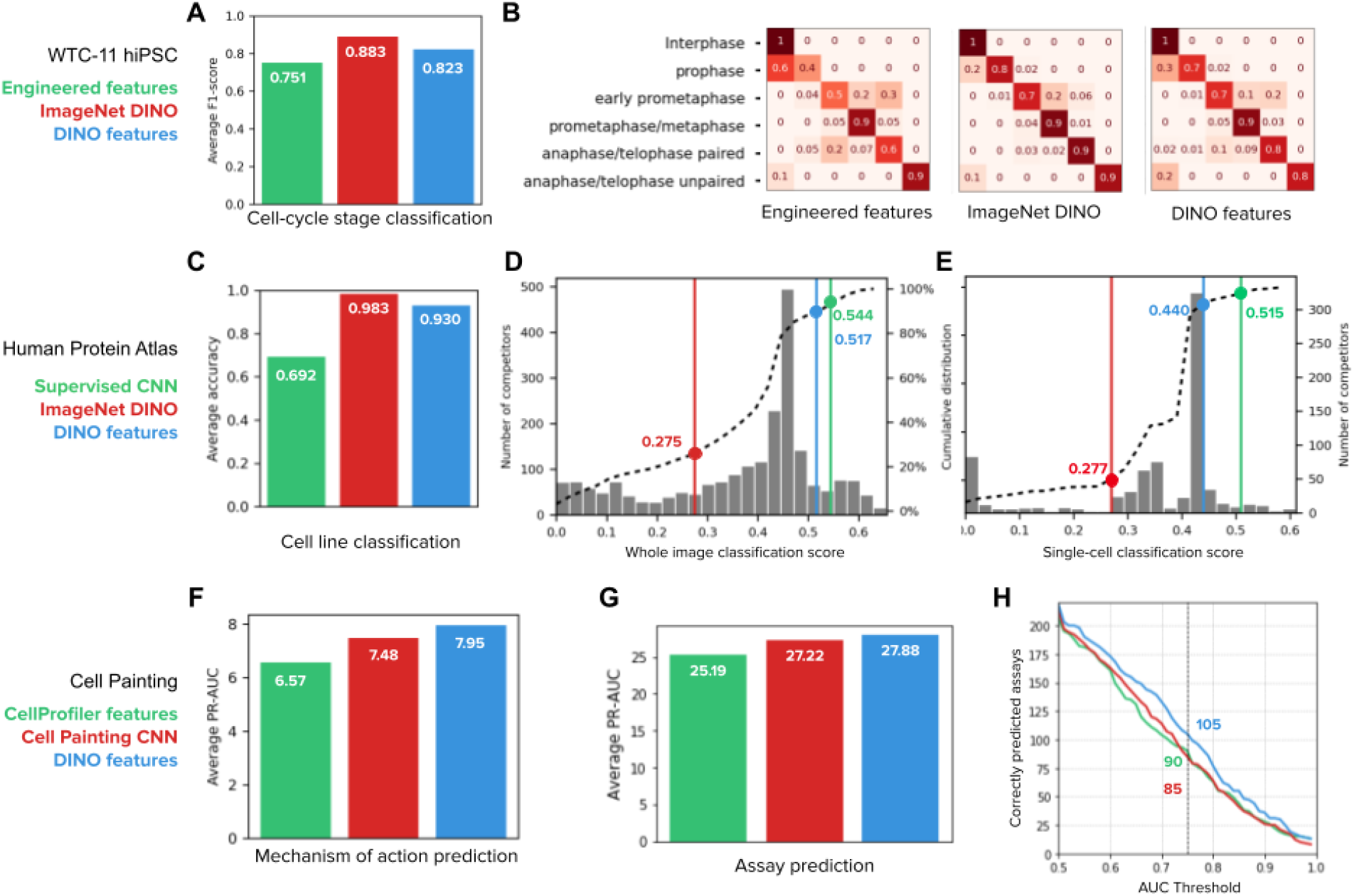
Quantitative evaluation of feature representations in supervised downstream tasks. A) Cell-cycle stage classification task in the Allen Institute WTC-11 hiPSC dataset. The bar plot shows the macro average F1-score over six classes. The three classifiers have the same multi-layer perceptron (MLP) architecture (Methods) and only differ in the input features: engineered features ^45^(green), DINO features trained on ImageNet (red), and DINO features trained on WTC11 images (blue). B) Confusion matrices of the three classification models in A, each showing the normalized true positive counts for the six cell-cycle stages (Methods). C) Cell line classification accuracy on whole-images of the Human Protein Atlas (HPA). The bar plot shows the average accuracy over 35 cell line classes for three identical MLP classifiers that differ in the input features: DINO features trained on ImageNet (red), the top performing CNN model (Bestfitting team) supervised for protein localization in the whole-image classification Kaggle competition ^21^ (green), and DINO features trained on HPA (blue). D) whole-image protein localization classification challenge ^21^. The histogram shows the distribution of the number of Kaggle competitors (left y-axis) according to their performance (x-axis, higher is better). Three classifiers trained with the input features in C are highlighted in the plot with lines and points of the corresponding colors. The dashed line is the cumulative distribution of the number of competitors (right y-axis). E) Single-cell protein localization classification challenge ^22^. The histogram shows the distribution of the number of Kaggle competitors (right y-axis) according to their performance (x-axis, higher is better). Three classifiers are highlighted in the plot as in D. The dashed line is the cumulative distribution of the number of competitors (left y-axis). F) Mechanism of action prediction for compounds in the LINCS Cell Painting dataset ^47^. Three identical MLP classifiers are trained with input features: CellProfiler features (green), Cell Painting CNN features ^29^(red), and DINO features trained on Cell Painting images (blue). G) Compound bioactivity prediction for small molecules in the BBBC036 Cell Painting dataset ^53^. The bar plot compares three classifiers as in F. The y-axis of the bar plot is the average PR-AUC for 270 assays. H) Number of assays correctly predicted by the three classifiers in G as a function of the AUC threshold. The highlighted numbers correspond to a AUC threshold of 0.75.

DINO features were also highly competitive in Kaggle-hosted, public benchmarks on both HPA whole-image (containing multiple cells) and single-cell datasets: an MLP classifier trained on frozen DINO features ranks in the top 13th and 12th percentile respectively, among hundreds of highly specialized submissions (Figure 3D,E). Using the same MLP classifier architecture, we further compared ImageNet DINO features, HPA DINO features trained on whole-images or single-cells, as well as features from the 1st and 2nd ranked solutions in the single-cell and whole-image classification challenges, respectively (Bestfitting^21, 22^ and DualHead^22^). Note that the top competitors designed ensemble models with many CNNs to improve their results ^21, 22^. To compare the quality of features rather than classification strategies, we evaluated a single CNN from their ensemble to compare with single DINO networks. The results confirm that DINO learns features that perform comparably with fully supervised CNNs in solving protein localization classification, exhibiting small performance differences (only 0.03 and 0.08 absolute points in the whole-image and single-cell challenges respectively, Figure 3D,E). Importantly, our approach drastically reduces the need for supervised training using an MLP with only 5.3x10^5^ parameters compared to 9.1x10^6^ parameters in a *single* network inside the ensemble of Bestfitting^21, 22^ (∼17x reduction, Figure S4E,F). With less supervised parameters, our approach may require less manual annotations too.

In addition to protein localization phenotypes, DINO features also encode cell-line variation efficiently. We compared three classifiers trained with ImageNet DINO features, features extracted from a network from the Bestfitting supervised CNN ensemble ^21, 22^, and HPA DINO features to test their ability to predict cell lines in a holdout set. As expected, DINO features succeed (0.93 classification accuracy) while the supervised CNN model underperforms (0.69 classification accuracy) because it was not optimized to account for factors of variation other than protein localization (Figure 3C for HPA whole-images, and Figure S4D for HPA single-cells). This training specialization limits the diversity of analyses that can be performed with images. For example, simultaneously identifying protein localizations and cell types through supervised approaches would require additional annotations that may not be readily available in other applications (e.g. complex tissue slides). In contrast, DINO delivers high-quality features over multiple factors of variation via a single, self-supervised learning objective.

### DINO improves the prediction of compound bioactivity

Imaging is extensively used to quantify the effects of pharmacological compounds at high throughput where the phenotypes of interest cannot be annotated by hand. The CPC dataset is an example of such an application. Here, existing knowledge from chemical biology is used to create predictive models of compound bioactivity by looking at the response of cells to perturbations in images ^54, 55^. We find that DINO improves the ability of such models to predict the MoAs of compounds and the results of diverse bioactivity assays using Cell Painting images^56^. For this evaluation, we first trained a DINO model with the same dataset as in the Cell Painting CNN-1 model ^29^ (Methods). Next, we computed features and created well-level representations for the LINCS dataset to predict MoAs ^47^, and for the BBBC036 dataset ^46^ to predict the readouts of 270 assays (see below).

For MoA prediction, we used annotations from the Connectivity Map project, which were featured in a Kaggle challenge to classify molecules based on transcriptional profiling ^57^. The evaluation protocol was adapted for image-based profiling using the LINCS Cell Painting dataset ^47^, a library of more than 1,500 compounds screened on A549 cells. We used the same profiling pipeline and classifier models (Methods) to compare three feature representations: engineered features obtained with CellProfiler ^58, 59^, features obtained with weakly supervised Cell Painting CNN-1 model ^29, 60^, and DINO features trained on Cell Painting images. The results show that learned features improve MoA classification compared to engineered features, and DINO features perform better than the CNN model (Figure 3F). Note that MoA classification is approached at the well-level instead of classifying images of single cells. The latter could potentially leverage single-cell heterogeneity to improve prediction performance even further.

We also evaluated the problem of predicting compound activity in the BBBC036 Cell Painting dataset with 30,000 small molecules screened on U2OS cells ^46^. The goal here is to use image-based features to determine if compounds are hits in one of 270 historical assays collected at the Broad Institute, given a sparse set of assay readouts ^53^. We followed the same profiling pipeline and classifier models (Methods) to compare CellProfiler features, Cell Painting CNN-1 features, and DINO features, following a well-level prediction approach. DINO features improved performance with respect to both CellProfiler and the CNN model, achieving higher overall precision-recall scores (Figure 3G, 10% and 2% improvement over baselines, respectively) and predicting bioactivity for more assays than the baseline models at all thresholds of accuracy (Figure 3H, 23% and 16% improvement over baselines, respectively).

Together with MoA prediction, these results confirm DINO’s ability to identify phenotypic variation across thousands of perturbations to approach hundreds of prediction tasks, holding great potential to advance high-throughput drug discovery projects.

### DINO enables unbiased profiling of cellular morphology

Unbiased profiling of biological conditions aims to uncover the relationships among groups of samples, to discover similarities or differences across them. DINO features are effective for unbiased image-based profiling, as indicated by its ability to cluster samples in a biologically meaningful manner (Figure 2). Here, we aggregated single-cell DINO features according to the conditions of interest in each dataset to build morphological profiles, then computed the cosine similarity across all conditions, and finally, clustered the similarity matrices to reveal groups related by their morphological structure (Figure 4).

**Figure 4.**
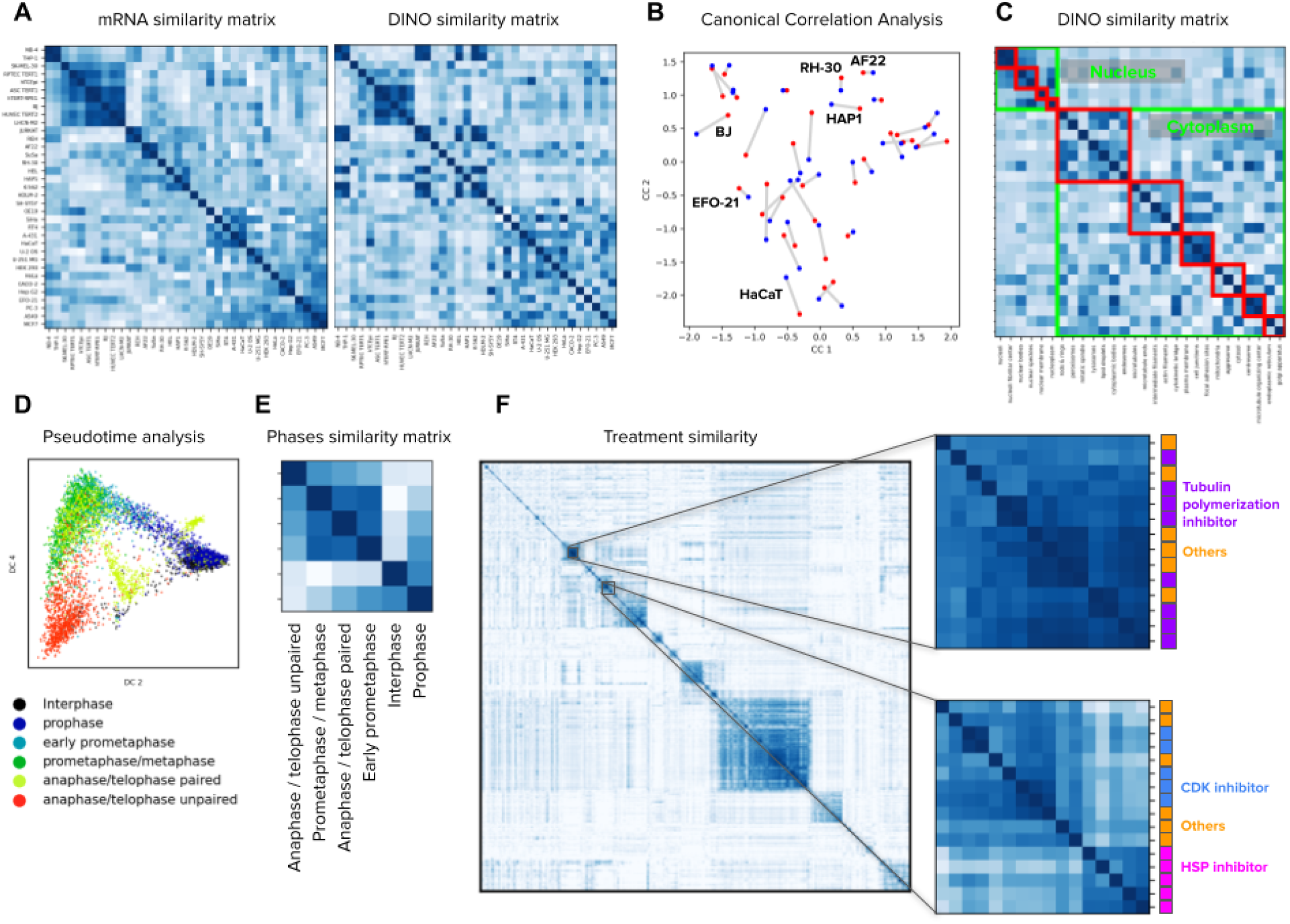
Image-based profiling of cellular state under different conditions. A) Cell line profiling in the Human Protein Atlas dataset using bulk mRNA levels (left) and aggregated DINO features (right). The matrices display the cosine similarity among cell lines (rows and columns) according to mRNA levels or imaging features. The order of rows and columns in both matrices follows groups determined by hierarchical clustering of the mRNA similarity values. B) Canonical correlation analysis between mRNA readouts and DINO features aggregated by cell line in the HPA dataset (Methods). Each point in the plot corresponds to one cell line. Red points are the mRNA representation and blue points are the DINO features representation. Lines between points indicate the correct connection between the two representations for one cell line. A representative subset of the cell line points are annotated. C) Matrix of cosine similarities among DINO features aggregated by protein localization groups, with rows and columns ordered by the ground truth annotations. The clusters highlighted in green are protein localizations in the nucleus or the cytoplasm. The red clusters correspond to secondary groupings of the protein localizations, annotated by experts^27^ (Methods). D) Pseudotime analysis of cell cycle stages in the WTC11 dataset using the Diffusion Pseudotime (DPT) algorithm ^63^ on the features extracted by DINO trained on ImageNet. Points in the plot are single cells colored by cell-cycle stages. E) Matrix of cosine similarities between DINO features aggregated by cell-cycle stage groups in the WTC11 dataset. F) Matrix of cosine similarities between DINO features aggregated at the treatment-level in the LINCS Cell Painting dataset. The two matrices in the right are zoomed-in views of two groups of compounds that share similar mechanism-of-action labels, indicated by the colors and names in the right.

The HPA dataset includes images of 35 human cell lines, which may be related to each other by e.g. anatomical origin or disease state. We aggregated DINO features to build cell-line profiles (Methods) and reveal their similarities (Figure 4A). For comparison, we created the cell-line similarity matrix according to transcriptional (RNAseq) profiles available through HPA ^61^ and used the Mantel similarity ^62^ to estimate their correlation (Methods). The RNAseq matrix has a Mantel statistic of 0.47 with the DINO matrix; higher than the statistic with the matrix of supervised CNN features (0.31) and the matrix of ImageNet DINO features (0.40). Canonical correlation analysis (CCA) also shows high correspondence between both cell line representations, obtaining 1NN accuracy of 0.25 for DINO features, which is 3 times better than 0.08 for supervised CNN features (Figure 4B, Methods). While the similarities between cell lines as captured by either RNAseq or cellular morphology are not expected to be identical, the results highlight how DINO features encode cell line variation consistent with biological relationships identified by orthogonal methods.

Aggregation of DINO features according to the 28 HPA protein localization groups in the HPA dataset and ordering them by the ground truth protein localization hierarchy annotated by experts ^27^ also results in a meaningful similarity matrix (Figure 4C), revealing protein localizations in the nucleus and in the cytoplasm as two major clusters, as well as several minor clusters within them (Methods). Importantly, these broadly recapitulate ground truth annotations, achieving a Mantel statistic between the DINO similarity matrix and the ground truth annotations of 0.52, outperforming the supervised CNN features (0.48). We conducted a similar profiling analysis for cell cycle stages in the WTC11 dataset and observed clustering in two major groups: interphase/prophase cells and mitotic cells (Figure 4E). Analysis of single-cell DINO features using the diffusion pseudotime (DST) algorithm ^63^ further revealed a cycle-like ordering of cells (Figure 4D) in a sample of images balanced to represent the six stages equally (Methods).

Finally, we created perturbation-level profiles for 1,249 compounds in the LINCS Cell Painting dataset by aggregating single-cell DINO features. Visual inspection of the cosine similarity matrix suggests these profiles naturally uncover similarities between perturbations that share similar MoA annotations (Figure 4F). To quantify the ability of DINO features to group compounds with similar MoAs, we followed the benchmark evaluation presented in the Cell Painting CNN-1 study ^29^, which is based on perturbation matching with nearest-neighbors search on three Cell Painting datasets. DINO features performed better than CellProfiler features in all three datasets (3%, 5% and 22% improvement in BBBC037, BBBC022 and BBBC036, respectively), and exhibited competitive performance with respect to the weakly supervised Cell Painting CNN features (Figure S6, ∼3% below in all datasets). Together, these results underpin the inherent biological fidelity of DINO features, and their potential to enable new and complement existing routes for unbiased biological discoveries.

### DINO captures single-cell heterogeneity

Imaging can naturally offer single-cell resolution, which allows to quantify cell state variations and heterogeneity for studying increasingly specific biological events. DINO features encode variations of cellular morphology that include the spatial organization of subcellular patterns, and illumination variations that correspond to levels of protein expression.

The HPA images were manually annotated in terms of single-cell variability, divided into spatial and intensity variance; spatial variance indicates that the same protein localizes to different subcellular structures, while intensity variance indicates different levels of protein expression. The HPA provides these annotations at the gene level, where different genes are labeled as either spatially varied, intensity varied, or both.

We found that DINO features are able to rank genes according to their heterogeneity score (Figure 5). We ranked genes according to the standard deviation of single-cell feature vectors, and then calculated the percentage of genes annotated as spatially or intensity heterogeneous within the top-k genes (Methods). The result shows that DINO features from a model trained on the HPA single-cell dataset encodes single-cell variation useful to correctly predict the ground truth heterogeneity of genes with peak accuracy of 64% at the top k=24 results, compared to 48% at k=34, and 32% at k=79 for a supervised protein localization CNN and an ImageNet pretrained DINO model, respectively (Figure 5B). The peak prediction improves to 70% when genes are ranked based on the variance of all single-cells that are stained for each gene (Methods).

**Figure 5.**
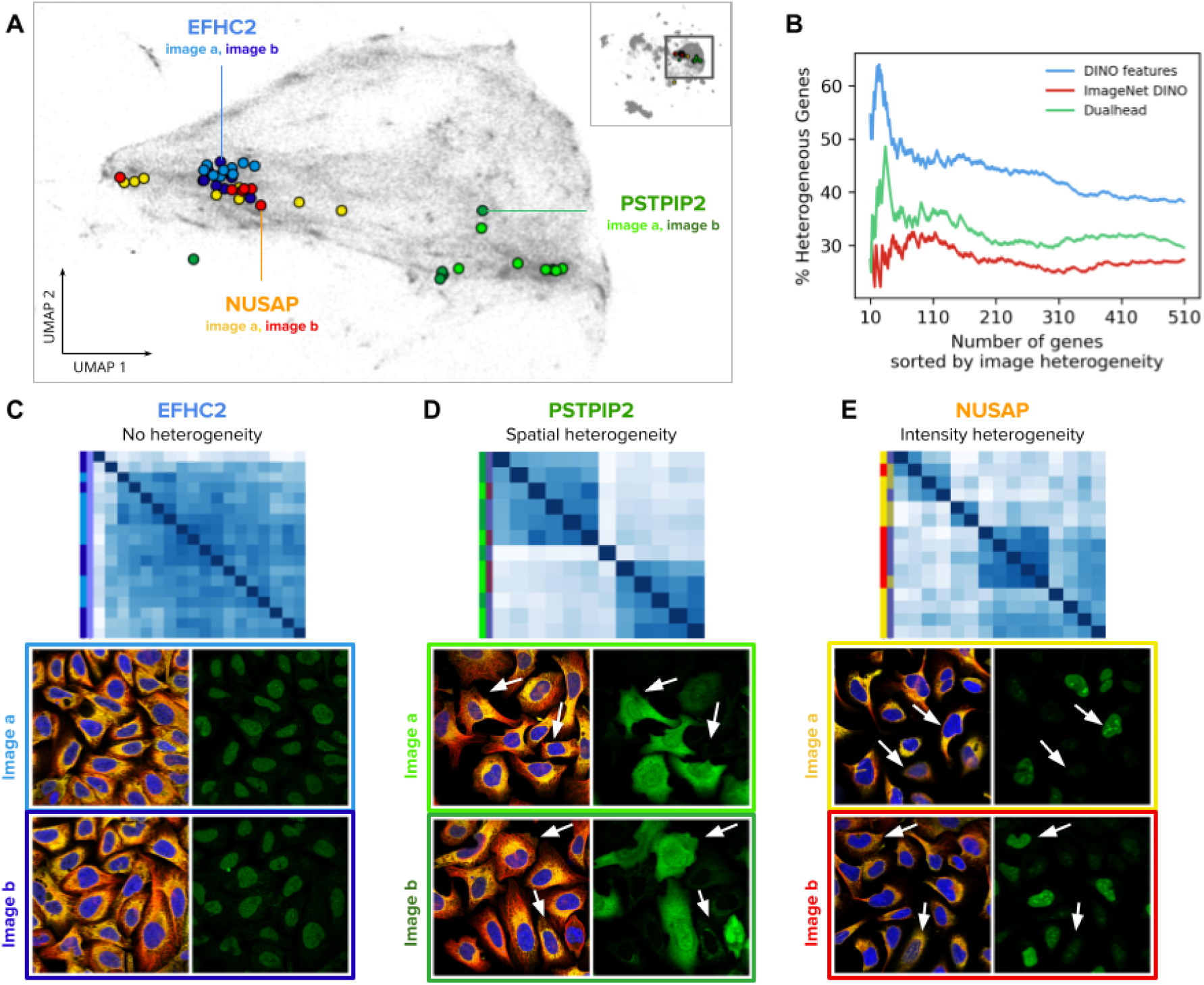
Single-cell heterogeneity of protein localization patterns in the Human Protein Atlas. A) UMAP visualization of single-cell DINO features. The small map in the top-right corner shows all single cells in the HPA dataset and highlights the cluster of U2OS cells, which are presented in the main plot. Single cells from two images of the genes EFHC2 (blue), NUSAP (orange), and PSTPIP2 (green) are displayed in colors to illustrate heterogeneity patterns. Panels C, D, and E use the same color convention of the genes and the images (shades of the corresponding gene colors). B) Ability of morphology features to capture single cell heterogeneity. The x-axis represents the ranking of genes according to the variance of single-cell features in an image (Methods). The y-axis represents the proportion of genes labeled as heterogeneous according to existing annotations in the HPA website. C, D, and E are examples of heterogeneous protein localization patterns, and show cosine similarity matrices of DINO features for single cells associated with three genes: EFHC2, PSTPIP2, and NUSAP, respectively. The images below the matrices are the source of the single cells in the analysis; left images: microtubules (red), endoplasmic reticulum (yellow), and nucleus (blue); right images: protein channel - arrows indicate cells exhibiting different protein expression patterns according to the type of heterogeneity. The color labels follow the conventions in A.

Our method of calculating single-cell heterogeneity within genes is better at predicting intensity variance than spatial variance. We found that single-cell DINO features show high variance for intensity heterogeneous genes, and a wider range of variances for heterogeneity in protein localization (Figure S5).

### Vision transformers encode biologically meaningful subcellular features

ViTs decompose images into a sequence of patches that are transformed using attention operations (Methods). We found that these transformed patches, also called patch tokens, capture meaningful information about subcellular structures. In contrast to CNNs, which pool local features into aggregated vectors, ViTs preserve the resolution of the input sequence in all the layers of the network, enabling unbiased analysis at subcellular resolution using individual patch tokens. We analyzed the emerging properties of the patch tokens and their corresponding attention maps in the WTC11, HPA, and Cell Painting datasets (Figure 6).

**Figure 6.**
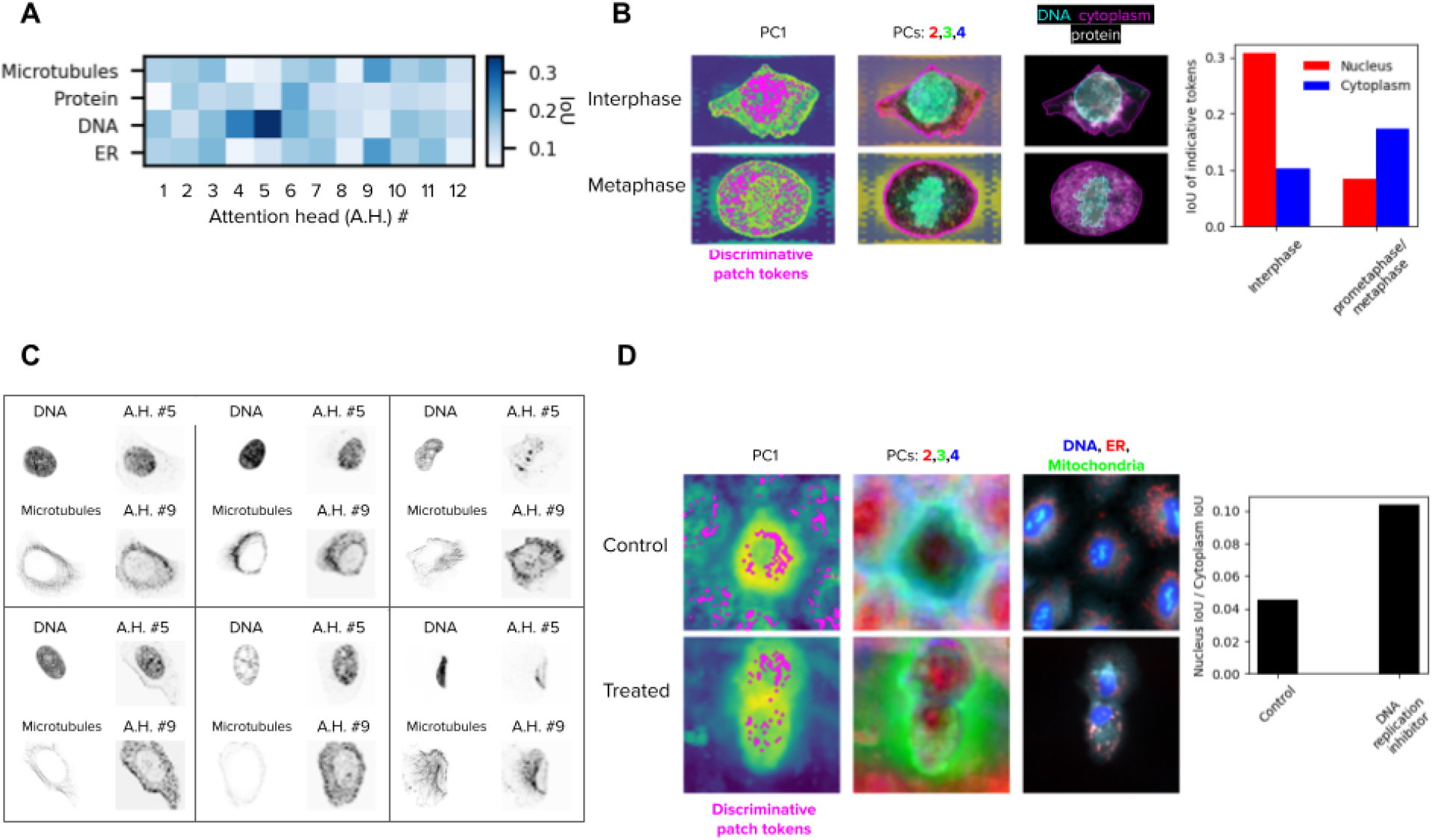
Vision transformers encode biologically meaningful features of subcellular structures. A) Matrix of the intersection-over-union (IoU) of fluorescent channels and attention heads in the last layer of the ViT DINO model trained on the HPA dataset. The IoU values are the average over 50 single cells. B) Left column: first principal component (PC) of the patch tokens for two cells, one of the interphase group and the other of the metaphase group in the WTC11 dataset. The locations of discriminative patch tokens are highlighted in magenta (Methods). Middle column: 2nd, 3rd, and 4th PCs of the key descriptors, portrayed in red, green and blue respectively. Right column: original single-cell image. Right: barplot of the average IoU of key descriptors and nucleus (red) or cytoplasm (blue) over the single cells of each group. C) Examples of fluorescent channels (left) and attention maps (right) of six single cells, where the attention head highly overlaps with the fluorescent channel according to A. D) Left: similar to B, but for cells from the DMSO control and a DNA replication inhibitor / STAT inhibitor in the LINCS dataset (compound ID: BRD-K35960502). Right: The average ratio of nucleus IoU over cytoplasm IoU per cell for each group. The treated group had a higher ratio, indicating relatively more discriminative patch tokens in the nucleus compared to the control group.

We looked at the twelve attention maps generated by the last layer of the ViT model trained on the HPA single-cell dataset and observed that they correlate with subcellular structures revealed by specific input fluorescent channels. To quantify this, we randomly sampled 50 single cells and computed the intersection-over-union (IoU) score between each attention map and each fluorescent channel of the input single-cell image to estimate their spatial overlap (Methods).

The resulting average interaction matrix (Figure 6A) shows how different heads attend to different subcellular structures. Some attention heads (e.g., #5, #9, Figure 6C) preferentially select individual fluorescent channels, while others (e.g., #2, #8) distribute attention equally across multiple cellular compartments. Interestingly, some heads pay attention to combinations of fluorescent channels, such as head #4, which simultaneously looks at the DNA and protein channels, potentially encoding meaningful subcellular interactions.

In addition, patch tokens can be used to highlight differences between groups of cells in a localized, high resolution manner. We randomly sampled 50 single cells from the interphase and 50 single cells from the metaphase cell-stages in the WTC11 dataset, and highlighted the patch tokens most useful for discriminating between the two groups (Methods, Figure 6B). These descriptors localize mostly in the nucleus for interphase cells, and mostly in the cytoplasm for metaphase cells, suggesting that features of those cellular compartments have the largest differences between the two cell-cycle stages. We found similar results when sampling 50 control cells and 50 treated cells (with a DNA replication inhibitor) from the LINCS dataset and highlighted the discriminative patch tokens. Visually, the main characteristic of this treatment seems to be cytokinetic failure, and the discriminative tokens suggest that our model captures the difference between control and treated cells as a reduced ratio between cytoplasmic to nuclear volume.

## Discussion

In this paper, we evaluated DINO as a general strategy for learning unbiased representations of cellular morphology. DINO is self-supervised and only requires microscopy images to reveal a wide range of relevant biological information for diverse downstream analyses. We used DINO to analyze datasets from three imaging studies that have different technical specifications as well as phenotypes of varying complexity, and evaluated performance in nine biological tasks, including cell-cycle staging, subcellular protein localization, and compound bioactivity prediction. These tasks involved morphological analysis of phenotypes at the population level, at single-cell resolution, and at subcellular resolution, highlighting the wealth of information that DINO can automatically discover from microscopy images.

Recent studies have investigated DINO and other self-supervised techniques for cellular profiling ^39, 64–66^, and histopathology analysis ^67, 68^, illustrating the diversity of specialized biological tasks that can be approached with representation learning for imaging. Our work extends prior studies in two major ways: first, we demonstrate that DINO captures multiple factors of variation in a hierarchical way, which can be used to address a variety of downstream analyses, making it an ideal tool for single-cell morphological profiling. This allows us to interrogate several aspects of the biology underlying imaging experiments like other single-cell omics; for example, using computational techniques from the single-cell genomics literature (e.g. diffusion pseudotime and Harmony). And second, while previous studies focused on a single, specialized task, ours systematically evaluate various imaging problems with different numbers of channels, technical variation, and biological relevance at different scales, demonstrating the general applicability of DINO for image-based analysis.

The field of self-supervision made significant strides in the past few years ^69–71^, and there have been extensive efforts to compare performance across techniques, including in bioimaging ^39^. DINO has consistently shown superior performance in these benchmarks, and continues to break records with improved scalability and performance quality ^72^. In our work, rather than comparing performance across different self-supervised techniques, we aimed to comprehensively investigate the emerging properties of vision transformers trained with DINO on biological imaging datasets, following insights observed in natural images ^30, 73^. The way DINO succeeds only by looking at images of cells is a remarkable example of how advances in artificial intelligence can power biological analysis. Imaging has long been considered a powerful tool for studying cellular phenotypes, but each experiment required specialized methods to extract useful image-based features. With advances in experimental protocols and the collection of ever larger imaging datasets, we expect DINO to make a significant contribution to future biological discovery.

## Acknowledgements

We thank A. Carpenter, S. Singh, M. Babadi, and M. Hirano for helpful discussions. This research is based on work partially supported by Muscular Dystrophy Association (Development Grant 628114), by the National Institutes of Health (award K99HG011488 to WMP), by the Broad Institute Schmidt Fellowship program (JCC), by the Broad Institute SPARC grants (JCC), and by the National Science Foundation (NSF-DBI award 2134695 to JCC).

## Methods

### Datasets

The first dataset we analyzed is the Allen Institute Cell Explorer. This dataset contains over 214,037 human induced pluripotent stem cells (hiPSC) from 25 isogenic cell lines, each cell line containing one fluorescently tagged protein via CRISPR/Cas9 gene editing. The cells are imaged in high resolution, 3D images, and are further stained with fluorescent biomarkers tagging the nucleus and the cell bodies. We used the max z-channel projection of these three fluorescent channels to train self-supervised feature extractors. Single cells were annotated by experts ^45^ with cell cycle stage labels, which we used to explore the biological relevance of feature representations.

The second dataset we analyzed is a subset of the Human Protein Atlas (HPA) Cell Atlas used for the Kaggle competition “Human Protein Atlas Image Classification”. The data is taken from 35 different cell lines with different morphology, such as A549 human lung adenocarcinoma cells or U2-OS human bone osteosarcoma epithelial cells. Each cell culture is treated with fluorescent dyes that bind to the nucleus, microtubules, endoplasmic reticulum, and a protein of interest, providing four image channels with different aspects of cellular morphology. The proteins of interest are divided into 28 classes of protein localizations, annotated by experts.

The original goal of the dataset was to test how well protein localization can be predicted given the four channel images. We also analyzed the HPA Cell Atlas single-cell subset, used for the Kaggle competition “Human Protein Atlas - Single Cell Classification”. This dataset is composed of 28 cell lines, and has the same fluorescent channels as in the HPA FOV classification challenge above, with the protein localizations being divided into 17 locations as well as another “negative” class.

We also analyzed a collection of Cell Painting datasets, including LINCS ^47^, BBBC036 ,^46^ and a combined single-cell resource for training ^29^. These datasets were obtained following the Cell Painting protocol ^56^, where cells are stained with six fluorescent dyes highlighting eight cellular compartments. Images are acquired in five-channels at 20X magnification. The LINCS dataset is a chemical screen of 1,249 FDA approved drugs in six concentrations, tested on A549 cells In our experiments we only use samples from DMSO negative controls and the maximum concentration of each compound. The BBBC036 dataset is a compound screen of 30,616 compounds at a single dose tested in U2OS cells. We use a subset of 16,170 compounds from this dataset for evaluation of compound bioactivity prediction. The combined Cell Painting dataset is a collection of subsets from five other Cell Painting datasets, covering gene overexpression perturbations, compound screens, two cell lines (A549 and U2OS), sampled to maximize technical, biological and phenotypic variation. We use this dataset for training feature extraction models.

### Cell Segmentation

In the WTC-11 dataset, we used the segmentation masks provided by the authors of the original study ^45^. These segmentations were obtained with the Allen Cell and Structure Segmenter ^74^, which segments 3D intracellular structures in fluorescence microscopy images using a combination of classic image segmentation and iterative deep learning workflows. In the HPA dataset, we used the HPA-Cell-Segmentation algorithm ^75^ recommended in the HPA single cell classification challenge to obtain images of single cells from the FOVs of the main dataset. The algorithm is based on pretrained U-Nets, and returns both cell segmentation and nucleus segmentation. In the HPA single cell DINO training, we trained the model on full images of single cells regardless of cell size, and inferred the features from center crops of size 512x512 pixels around the center of each single cell. Segmentation of Cell Painting images was performed with CellProfiler using a two steps approach: 1) The Hoechst (DNA) channel is segmented with global Otsu thresholding, and used as a prior for the next step. 2) Cell body segmentation with the watershed method in the ER or RNA channel.

### DINO algorithm

DINO (self-DIstillation with NO labels) is a self-supervised training algorithm that trains an unbiased feature extractor. DINO is composed of two networks, student and teacher, that share their architecture. Both networks receive an image and output a feature vector that is then passed on to a projection network that classifies the feature vectors to a logit vector of 65,536 values. The weights of the student network are trained with the cross entropy loss between the outputs of the student and teacher networks. The weights of the teacher network are updated using the exponential moving average of the student network. After training, the projection head is removed from the feature extractor networks, and the trained teacher network is preserved for feature extraction. The images that the two networks receive are views of the same sample, and augmentations are used to indicate which information is not important. Thus, as the two networks are trained to output a similar feature vector for all views of the same image, the feature extractor learns to be invariant to the changes between the views, and encode only the non-changing aspects of the image.

An important part of the DINO framework is the generation of image views using augmentations. Unlike feature engineering, where researchers introduce prior knowledge by explicitly calculating the aspects of the data that are believed to be important, in DINO, image augmentations aim to make the model invariant to aspects of the data that are believed to be irrelevant. Thus, by choosing specific augmentations for biological images, we can train DINO to learn important features, and to ignore irrelevant information. We used cross validation with a subset of 5,000 FOV images from the HPA dataset to test the effect of introducing different augmentations and trained DINO models for 100 epochs. A linear classification model was then trained on the resulting learned features to evaluate performance. We found the following augmentations to be unnecessary for images of cells: Gaussian blurring, solarization, and greyscale transformations. In addition, we found several other augmentations to be useful: randomly rescaling the intensity of the protein channel (in HPA), dropping the content of a random channel by zeroing out all pixels, replacing color jitter with random brightness and contrast changes to each channel individually, and distorting the image with artificial warping. The number of channels of the ViT networks is adjusted independently for each dataset to 3, 4 and 5 channels in the WTC11, HPA, and Cell Painting datasets, respectively.

### Vision Transformer (ViT) architecture

The architecture of the student and teacher networks can be any neural network that receives an image and outputs a feature vector. However, DINO shows its strength best with the vision transformer (ViT) ^35^. The ViT is based on the transformer architecture ^76^, which is built as a series of connected attention heads. Each attention head receives a list of n tokens, transforms them into keys, queries and values, and multiplies them in an attention head to get a nxn attention matrix. While the original transformer was used for natural language processing, where each token is a word, the vision transformer works with images, where each token is the encoding of an image patch of pxp pixels. One of the benefits of using ViT instead of convolutional neural networks is that the ViT keeps the resolution the same regardless of the depth of the network, allowing us to iteratively calculate more complex features while keeping the same patch-wise resolution.

We used one ViT model for each dataset. For the WTC11 dataset we used a ViT-base architecture with a feature vector of size 768, patch size of 8 pixels and 3 input channels. For the HPA FOV images dataset we used a ViT-base architecture with a feature vector of size 768, patch size of 8 pixels and 4 input channels. For the HPA single cell dataset we used a ViT-base architecture with a feature vector of size 768, patch size of 16 pixels and 4 input channels. For the Cell Painting dataset we used two models: For the supervised tasks, we used a ViT-small architecture with a feature vector of size 384, patch size of 16 pixels and 5 input channels. For the unsupervised task, we used a ViT-base architecture with a feature vector of size 768, patch size of 16 pixels and 5 input channels. When using the DINO models trained on ImageNet, we adapted the 3 channel model to the varied channel data by duplicating each channel 3 times to be an RGB image and extracting features from the resulting 3 channel image, then concatenating the features of all data channels.

### Data post-processing and visualization

In the WTC-11 dataset, we first reduced feature dimensionality by taking the top 100 principal components (PCs), retaining 91% of the variance in the data. Next, we used unsupervised UMAP to project features in 2D for cluster visualization (Figure 2A). Additional post processing was used to integrate cell lines and generate a different visualization. To correct for cell line variation, we used Harmony on the reduced features and projected the corrected embeddings in 2D using UMAP (Figure 2B). We used the cuML version of UMAP with the default parameters, except for spread (4), n_epochs (2600), transform_queue_size (20), and random_state (42).

In the HPA single-cell data, we also reduced feature dimensionality to the top 100 PCs, retaining 77% of the variance in the data, and then projected all data points in 2D using UMAP. We train the UMAP projection using image-level data points to smooth out technical variation, by averaging single-cell feature vectors from the same FOV, while taking only single-cells with a single protein localization, to prevent multi-label average noise. With this image-level UMAP model, we then projected all individual single-cell feature vectors (Figure 2D). In the protein localization case, we colored all samples with multiple labels with gray, and colored the samples with a single label with its appropriate color. This strategy of training UMAP with image-level averaged features followed by single-cell feature projection was necessary to correct the visualization of raw features, which is dominated by image-level similarities (Figure S4). In addition, we integrated cell-line clusters using Harmony to visualize protein localization variation (Figure 2). When visualizing the FOV image-level samples, we did not reduce the dimensionality, and applied the UMAP and Harmony directly on the raw features.

In the Cell Painting dataset, we sampled 10% of the 8M single cells of the training dataset to create a UMAP visualization without any transformations (Figure S1). We also computed single-cell features on the LINCS dataset, which was used for downstream analysis, and aggregated them at the well level. Next, we applied the sphering batch-effect correction method and trained a UMAP model to visualize mechanism-of-action clusters (Figure S1).

### Mutual information analysis for factors of variation

We calculate the mutual information between image features and existing image annotations (herein called factors of variation) to obtain a quantitative estimate of the type of information that feature representations encode ^49^. We followed this procedure: a discrete joint-distribution of a factor of variation and features is first calculated. The number of bins for the factors of variation distribution was set to the minimal number of unique values across all factors of variation for each dataset. In this way, we keep the number of bins constant and allow comparison among models and factors of variation. Next, we calculate the mutual information between factors of variation and individual features. The result is an estimation of how informative each feature is to each factor of variation across the dataset (Figure S3). After that, we rank features based on the factor of variation they are most informative for. Finally, we calculate the percentage of features that are most informative for each factor of variation (Figure 2C,F).

In the WTC-11 analysis, we considered three factors of variation: cellular structures, cell cycle stage annotations, and wells. The number of bins was set to 6 for all factors of variation. In the HPA analysis, we used images from the website dataset (not the kaggle dataset), since these included cell line annotations. We considered three factors of variation: cell lines, protein localization groups, and wells. The number of bins was set to 25 for all factors of variation. The full report of factor of variation analysis can be found in Figure S3.

### MLP classifier architecture

In all of our experiments involving supervised learning in the WTC11 and HPA datasets, we used a multi-layer perceptron (MLP) classifier that takes features as input and produces classification predictions as an output. The MLP classifier consists of three layers: 1) Input linear layer with 512 hidden units, ReLU nonlinearity, and dropout regularization with probability 0.5. 2) Hidden linear layer with 256 hidden units, ReLU nonlinearity, dropout regularization with probability 0.5. 3) Linear classification layer with as many output units as the number of classes for the specific problem. The classifiers were trained with the softmax loss function using a cosine learning rate schedule (initial rate of 0.01) with a varying number of epochs contingent on the dataset (see below).

### Cell cycle classification task in the WTC11 dataset

The cell cycle classification task in the WTC11 dataset consists of classifying each single cell into one of six mitosis stages annotated by experts (Interphase, prophase, early prometaphase, prometaphase/metaphase, anaphase/telophase paired and anaphase/telophase unpaired). To classify cell-cycle stages based on cell images, we first divided the data into training and test sets with 80 and 20 percent respectively. Next, we balanced the training data, as 95% of the samples in the full dataset belongs to one class: we resampled the training set to have 28,538 samples per stage (171,228 samples overall) by oversampling the less frequent stages and undersampling the more frequent stages. We trained an MLP classifier on the features extracted by the DINO models trained on the WTC11 data and on the ImageNet data, reporting the classification score on the unbalanced test set. For comparison, we trained an XGBoost model on the balanced dataset using the engineered features of the original WTC11 dataset.

### Cell line classification - Human Protein Atlas

For the cell line classification, we first extracted features using the frozen DINO model that was trained on the HPA data in a self-supervised fashion. We removed all samples that were taken from the Kaggle dataset, as they had no cell line label, and kept the samples from the HPA website. We used these features as inputs to an MLP classifier head. Since the cell line distribution is highly unbalanced and multi-labeled, we balanced the data prior to training: we resampled each image with a probability inversely proportional to the frequency of its label, such that samples with infrequent classes appeared more often. We scaled the features to have zero mean and unit standard deviation by calculating the mean and standard deviation of the raw data in the training set, and normalized the validation and testing sets using the same training set mean and standard deviation.

For FOV classification, we trained the classifier for 100 epochs. For single-cell classification, we trained the classifier for 10 epochs, accounting for the larger number of samples in the single-cell case (Figure S4). We used a binary cross-entropy loss with logits. We report the performance using macro averaged F1 score.

### Protein localization classification - Human Protein Atlas

For the protein localization classification, we first extracted features using the frozen DINO model that was trained on the HPA data in a self-supervised fashion. We used these features as inputs to an MLP classifier head. Since the protein localization distribution is highly unbalanced and multi-labeled, we balanced the data prior to training: we resampled each image with a probability inversely proportional to its least frequent label, such that samples with infrequent classes appeared more often. We scaled the features to have zero mean and unit standard deviation by calculating the mean and standard deviation of the raw data in the training set, and normalized the validation and testing sets using the same training set mean and standard deviation.

We trained the classifier with a scheduler following cosine annealing ^77^, an AdamW optimizer ^78^ with weight decay of 0.04, beta1 of 0.9, beta2 of 0.999, learning rate of 1e-4, and batch size of 512. For the FOV classification, we trained the classifier for 10 epochs. For the single-cell classification, we trained the classifier for one epoch, accounting for the larger number of samples in the single-cell case. We used a binary cross-entropy loss with logits, to train the classifier to provide a separate probability for each class in the multi-label task. We report the performance using macro averaged F1 score.

### Mechanism of action prediction with Cell Painting

We performed supervised mechanism-of-action (MoA) prediction on well-level features extracted with CellProfiler, Cell Painting CNN, and DINO models. The classification pipeline is adapted from previous work ^47^. Specifically, we trained an MLP to take different types of features as input and predict the MoA of the compound that was used to treat each well. Compounds can have multiple MoA annotations.

We used CellProfiler, Cell Painting CNN, and DINO features as the input to classifiers to predict MoAs for the same set of compounds and wells. For CellProfiler features, we used the publicly available features previously collected and preprocessed ^47^. For Cell Painting CNN features, we used DeepProfiler to extract features from pre-trained weakly-supervised EfficientNet models ^29^ . For all feature sets, we selected only the wells treated with the maximum dose of each compound and preprocessed to align their MOA annotations. As part of the original classification pipeline, we performed principal component analysis (PCA) on the features to get 25 principal components and concatenated them to the existing feature vectors. We then normalized all the features using z-score normalization. Compound stratification and train-test split is performed as described in the original benchmark to ensure zero overlap between the training and testing compounds ^47^.

For the classifier, we compared the performance of the top models for multi-label MoA prediction from the MoA Kaggle competition ^57^. We chose the best performing model which has a Residual Neural Network (ResNet) architecture and includes six fully-connected layers with batch normalization and drop out layers. During training, we performed five-fold cross validation for hyperparameter tuning. Since each compound has only a few replicates, we placed all wells treated with the same compound into the same fold, resulting in a leave-compound-out evaluation. The output of the classifier is a probabilistic value between 0 and 1 for each MoA label. We then evaluated performance by calculating the area under the precision-recall curve (PR-AUC) for each MoA and taking the micro-average of the results. Positive controls are excluded when calculating the PR-AUC score as they have many replicates and induced distinct outlier phenotypes.

### Compound bioactivity prediction with Cell Painting

In the downstream compound bioactivity prediction task, we used a dataset of historic assay readouts from the Broad Institute, compiled for multi-modal prediction evaluation ^53^. We focused our evaluation on morphology representations only, and compared CellProfiler features, Cell Painting CNN features ^29^, and DINO features. All features were extracted at the single-cell level from Cell Painting images of the BBBC036 dataset ^46^ for a subset of 16,170 compounds that have partial assay readout annotations in 270 assays. Features were aggregated at the well-level and then batch-corrected using the sphering transform to create treatment-level profiles for machine learning analysis. The learning problem consists of predicting the status of compounds (hit vs no-hit) according to the 270 assays given their phenotypic profiles and a very sparse matrix of assay results.

We train a predictive model for each feature type using Chemprop ^79^, taking treatment-level profiles as input. For training purposes, Chemprop also expects an assay-compound matrix (16,170 compounds x 270 assays) with binary values (hit or no hit) or empty values (not tested / unknown). The evaluation setup follows a scaffold-based partitioning of compounds to conduct a five-fold cross-validation. When phenotypic features are provided, a Chemprop model is an MLP with up to 6 hidden layers trained with a multi-task loss, where each assay is one task. At inference time, this MLP produces the hit probability for each assay-compound pair in the test set, which we use to calculate ROC-AUC for all assays. Finally, we summarize the ROC-AUC scores for each assay using the median over cross-validation splits and count the number of assays with median ROC-AUC > 0.9.

### Unsupervised MoA prediction with Cell Painting

In the downstream unsupervised MoA prediction task, we applied a nearest-neighbor classifier to find how useful the features are for finding the MoA of new queries. To quantify this, we used the folds of enrichment and the mean average precision methods used and described in depth in the baselines ^29^. Folds of enrichment calculates the odds ratio of a one-sided Fisher’s exact test, and the final quantity is the average over all query treatments. Mean average precision calculates the average precision for each query treatment.

### Similarity of aggregated profiles of cell populations

To create similarity matrices of cell lines, we first aggregate the whole image DINO features into cell population profiles by averaging all vectors of the same population (e.g., averaging the vectors of all FOVs from the same cell line). Next, we reduce the dimensionality of the aggregated features using PCA and preserve the top 10 principal components, keeping 73% of the variance. Then, we calculate the cosine similarity across all reduced aggregated profiles to obtain the similarity matrix in Figure 4A right. To create the similarity matrix of the bulk RNASeq data, we followed the same process, while reducing the dimensionality of the Transcripts Per Million (TPM) for all genes across cell lines to the top 10 principal components, resulting in a 10-dimensional gene expression representation per cell line. For the order of the rows and columns, we calculated the hierarchical clustering of the RNASeq data, and ordered both the DINO and the RNASeq similarity matrices accordingly. To quantitatively compare the two similarity matrices, we used the Mantel test ^62^.

To further compare between the representations of the DINO features and the RNASeq data, we used the Canonical Correlation Analysis (CCA) ^80^ to project both data modalities in a common subspace spanned by the top 2 directions of maximal correlation. In this subspace, we calculated the Euclidean distance among cell lines across modalities and identified the top-1 nearest neighbor for each cell line. We used accuracy to estimate the correctness of cross-modal matches.

To create the similarity matrices between protein localization groups, we first removed the samples that have multiple protein localization labels, and then reduced dimensionality by taking the first 10 principal components and then aggregating with average, similar to the cell lines above. Next, we calculated the cosine similarity among all protein localization profiles to create the similarity matrix. We order the rows and columns based on existing annotations of the hierarchical organization of protein localizations ^27^. According to the annotations, the protein localization groups are divided by the superclasses nuclear and cytoplasmic. The nuclear superclass is subdivided into nucleus, subnuclear structures, nucleoli, and nuclear membrane. The cytoplasmic superclass is subdivided into cytoplasm, cytoskeleton, MTOC, secretory, and cell periphery.

To create the similarity matrix for perturbations in the LINCS Cell Painting dataset, we aggregated profiles by averaging single-cell features from the same perturbation, and then used the cosine similarity to create the matrix in Figure 4F. Due to the sparsity of the similarity matrix, we kept the top 50% of the treatments, based on their average similarity to other treatments. We ordered the rows and columns using hierarchical clustering.

### Pseudotime analysis

For the pseudotime analysis of cell morphology in the WTC11 dataset, we used the diffusion pseudotime (DST) algorithm ^63^, which estimates a pseudotime value by calculating the distance from a core origin sample to other samples in a neighbor graph. We first reduced the dimensionality of DINO features by taking the top 50 PCs, then balanced the dataset by taking 914 samples from each of the six cell-cycle stage classes. Finally, we ran DST using the scanpy Python package ^81^. To visualize the pseudotime result, we plotted the second and third components of the diffusion matrix for all samples, coloring them by the ground truth cell stage labels in Figure 4D.

### Single cell heterogeneity

To calculate the single-cell heterogeneity of the subcellular localization of proteins in the HPA dataset, we first grouped single-cell features from the same FOV, and calculated the standard deviation of the features. Next, we ranked genes by taking the maximum standard deviation among the FOVs stained for their corresponding protein. This value was used as the heterogeneity rank of the gene. To compare gene rankings across feature spaces, we used gene variability annotations available in the HPA website, and counted the number of genes annotated as varied in the top k genes of the ranked list. The annotations include three cases of variability: spatial variability, intensity variability, and not variable; many genes may have missing annotation. For the selected examples, we analyze each gene by sampling one well, and plotting the single cells from the two available FOVs in that well, and report one example for each of the three types of annotations.

### Evaluation of subcellular features

To quantify how DINO attends to the input fluorescent channels in the HPA dataset, we calculate the Intersection over Union (IoU) between the attention maps and the individual channels. We first binarized the fluorescent channels by applying Otsu’s thresholding method ^82^. We binarized the attention maps in the same way, by taking the output of the attention heads in the last layer with respect to the [CLS] token and applying Otsu’s thresholding. We calculated the IoU between each attention head and each channel, which results in a channel-attention interaction matrix (Figure 6A).

To identify differences in subcellular features between experimental groups, we sampled 50 DMSO control cells and 50 cells treated with a DNA replication / STAT inhibitor from the LINCS Cell Painting dataset (compound ID: BRD-K35960502). We extracted key descriptors from the last layer of the transformer network using dense, overlapping tokens ^83^ , and removed all background patch tokens that are not inside the cell mask. Next, we reduced the dimensionality of the patch tokens to the top 4 PCs, and checked how useful they are to classify cells into control vs treated conditions. A kNN classifier was used on the combined set of tokens from all single cells to identify those surrounded by other tokens of the same group (control vs. treated) and not by tokens of the same exact cell (such that the chosen tokens will highlight aspects of the category instead of the individual sample). Specifically, this was done by choosing the tokens that had 5 neighbors of the same group, but no neighbors of the same cells within the top 2 neighbors. We created a new image mask using the location of these patch tokens, which we call discriminative patch tokens. Finally, we quantified how many of the tokens are located in the cytoplasm vs the nucleoplasm to estimate the relative ratio of cellular compartment localization (Figure 6D). We followed the same procedure with the WTC11 data, by sampling 50 cells in the interphase stage and 50 in the metaphase stage, both from cell lines highlighting the Mitochondria cell structure (Figure 6B).

## Code availability

We share all code necessary for training the DINO models on the three datasets in this repository: https://github.com/broadinstitute/DINO4Cells_code

We share all code necessary for reproducing the results, including classification, analysis and figure generation, in this repository: https://github.com/broadinstitute/Dino4Cells_analysis

## Notes

### Competing Interest Statement

The authors have declared no competing interest.

