## Supplemental figures for "Unbiased single-cell morphology with self-supervised vision transformers"

### Supplementary Material

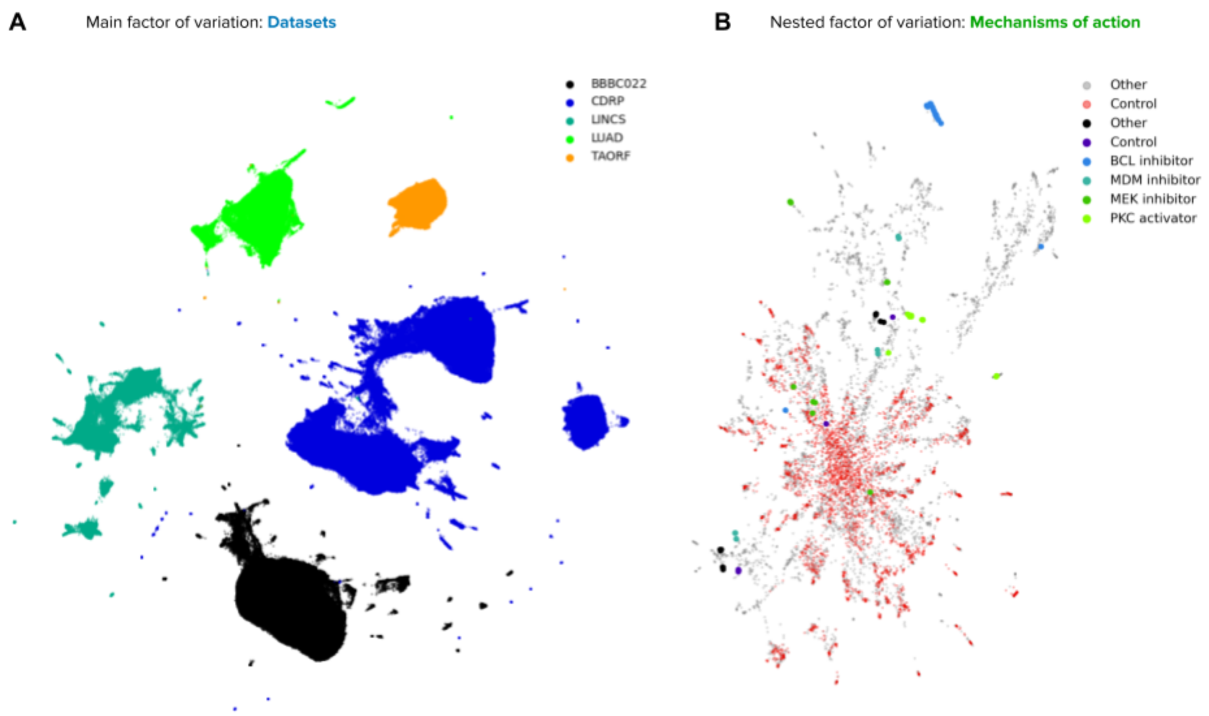


***Supplementary Figure S1. Visualizations of the Cell Painting morphology feature space automatically discovered by DINO.*** *A) UMAP visualization of single-cell morphology features obtained with DINO for images in the Cell Painting dataset. The plot shows 812,560 single cells, which are 10% of the data, and is colored by the source dataset. B) UMAP visualization of well-aggregated features obtained with DINO for images in the LINCS dataset, after correction for batch effects with the sphering transformation. Red points are DMSO wells, non-grey points indicate perturbations from selected mechanisms of action, and gray points are other wells.*

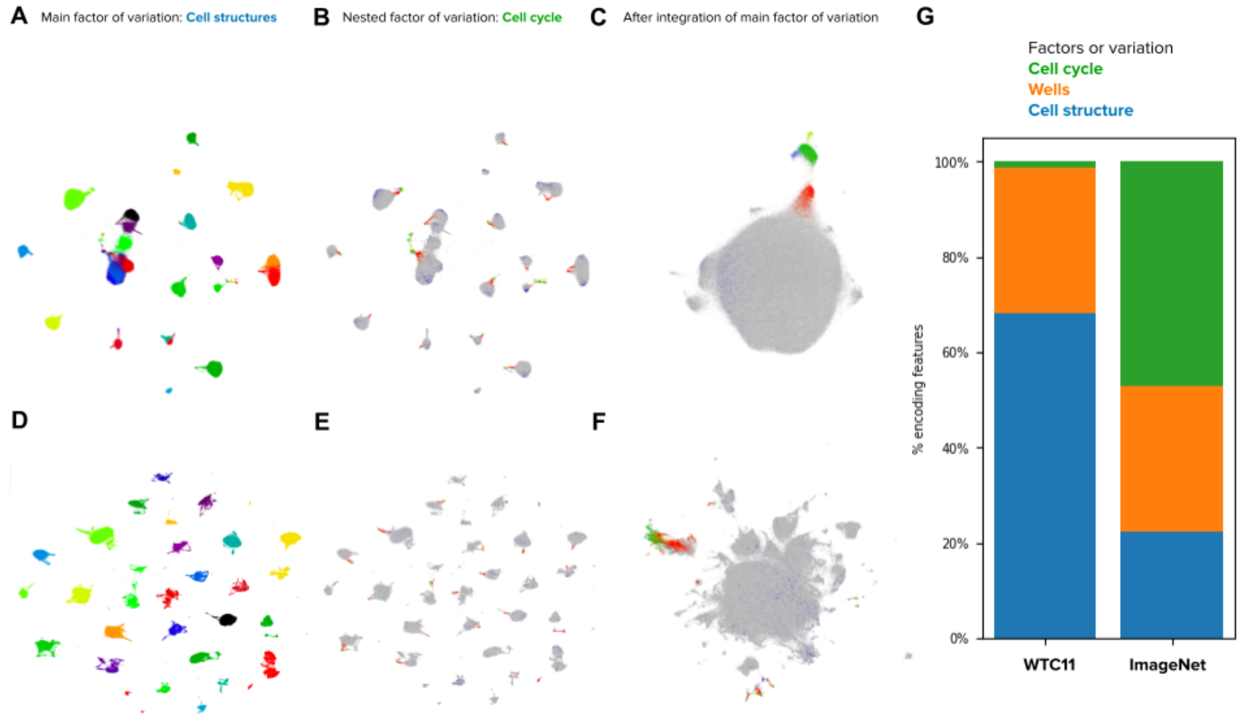


***Supplementary Figure S2. Comparison between the feature spaces discovered by DINO pretrained on ImageNet and DINO trained on the WTC11 dataset.*** *A) UMAP visualization of single-cell morphology features obtained with DINO for images in the WTC11 dataset. The plot shows the samples colored by cell structures. The features were obtained using DINO pretrained on ImageNet, without further training on WTC11. B) Same as in A, colored by the cell-cycle stage. C) ImageNet DINO features of WTC11 cells , integrated with Harmony over cellular structure groups. Points are colored by the cell-cycle stage as in B. D) Single-cell features obtained with a DINO model trained on WTC11. E) Same as in D, colored by the cell-cycle stage. F) DINO features trained on WTC11 cells and integrated with Harmony over cellular structure groups. Points are colored by the cell -cycle stage as in E. G) Stacked-bar plot displaying the fraction of features that are strongly associated with three factors of variation annotated in the WTC-11 hiPCS dataset, for features extracted using DINO pretrained on ImageNet and for features extracted using DINO trained on WTC11.*

Supplementary Figure S2 compares the feature spaces extracted using two DINO models, one pretrained on ImageNet and the other trained on WTC11. The feature space extracted using the pretrained model (Figure S2A,B and C) shows a smoother feature space than the one extracted using the model trained on WTC11 (Figure S2D,E and F). This is most evident when looking at the feature space UMAP after using Harmonize to remove the cell structure information, as the trained DINO feature space contains more clusters than the pretrained UMAP. This receives further support by quantitatively comparing the amount of features encoding the factors of variation in Figure S2G: While the features extracted using the pretrained model encode all factors of variation, the features extracted by the trained model encode the two factors of variation prone to technical variation (wells and cell structure) in more of the feature space. We belief this may be due to the self-supervised model learning all variation in the data, including technical variation, while the pretrained model does not have the opportunity to do so.

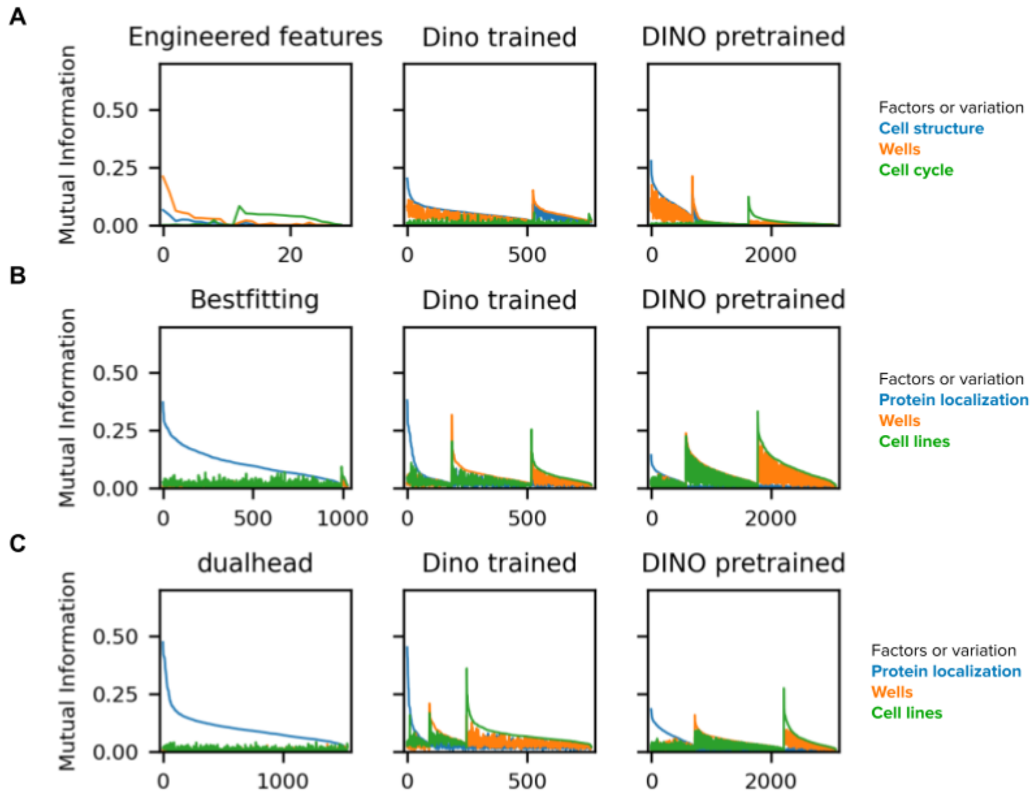


***Supplementary Figure S3. Mutual information between individual features and factors of variation.*** *A) Each color line represents the amount of mutual information (y-axis) between each feature of the WTC11 dataset (x-axis) and one factor of variation. The features are ordered by the amount of mutual information between them and three factors of variation: first the features with the dominant factor of variation being the cell structure, then the wells, and finally the cell cycle. We compare three feature extraction models: the engineered features, DINO trained on WTC11, and DINO pretrained on ImageNet. B) Same as in A, but for the HPA FOV dataset, with the factors of variation being protein localization, wells and cell lines. The compared feature extraction models are the Bestfitting Kaggle group, DINO trained on the HPA FOV dataset, and DINO pretrained on ImageNet. C) Same as in B, but for the HPA single-cell dataset. The compared feature extraction models are the dualhead Kaggle group, DINO trained on the HPA single-cell dataset, and DINO pretrained on ImageNet.*

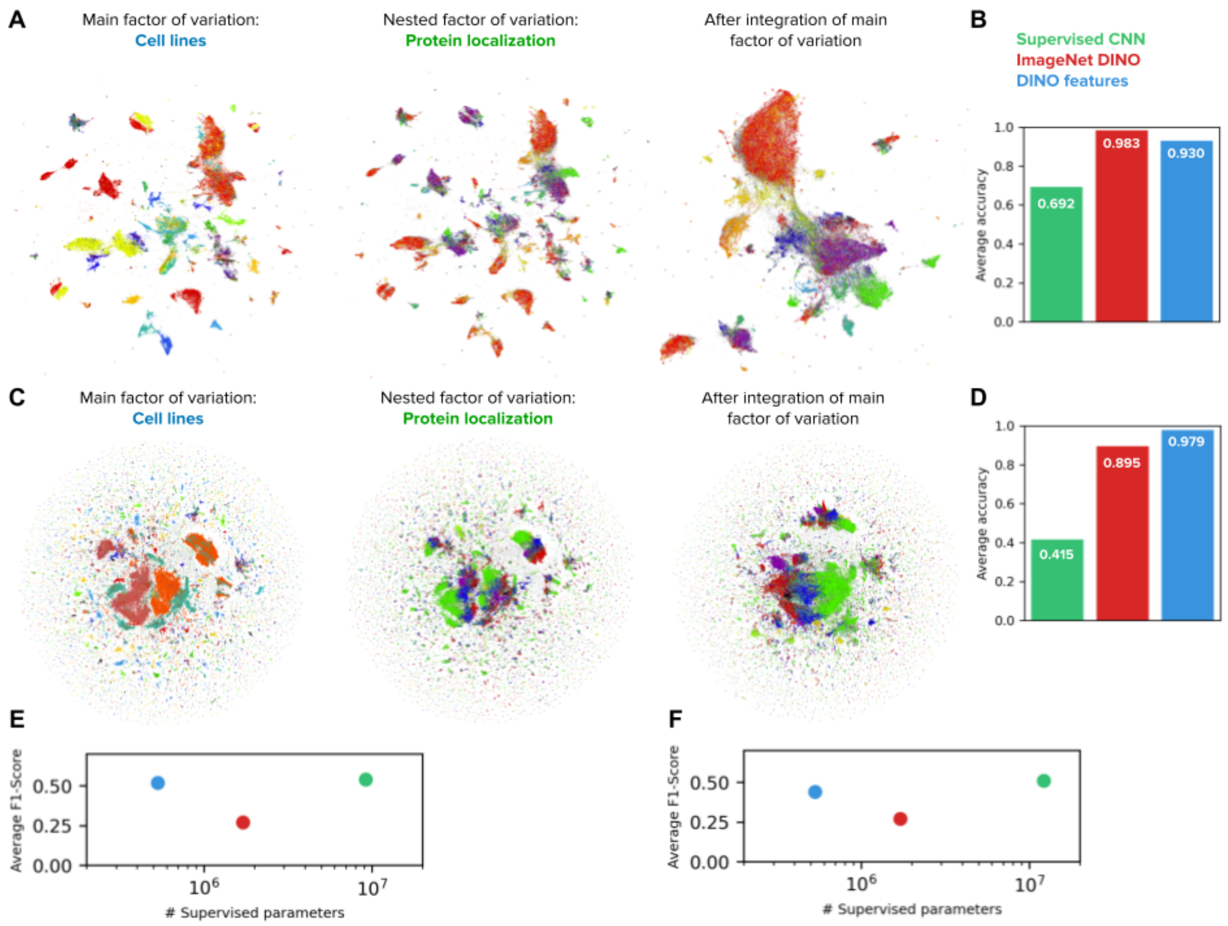


***Supplementary Figure S4. Visualizations of the feature spaces discovered by DINO trained on the HPA FOV dataset and DINO trained on the single-cell dataset.*** *A) UMAP visualizations of FOV morphology features obtained with DINO for images in the HPA FOV images dataset. The plot in the left shows the samples colored by cell lines, and the plot on the middle shows the samples colored by protein localizations. The plot on the right shows the same samples integrated using Harmony over the cell lines, and colored by protein localizations. B) Cell line classification accuracy on FOV images of the Human Protein Atlas (HPA). The bar plot shows the average accuracy over 35 cell line classes for three identical MLP classifiers that differ in the input features: DINO features trained on ImageNet (red), the top performing CNN model (Bestfitting team) supervised for protein localization in the FOV classification Kaggle competition [21] (green), and DINO features trained on HPA (blue). C) Same as in A, with features extracted from the HPA single-cell dataset using DINO trained on the HPA single-cell dataset. Unlike Figure 2, the feature vectors were not aggregated before training the UMAP. The raw features are dominated by technical variation, creating the isolated clusters seen in the UMAPs embeddings. D) Same as in B, for the HPA single cell dataset. The models compared are the DINO features trained on ImageNet (red), the 2nd best performing CNN model (dualhead team) supervised for protein localization in the single cell kaggle competition (green), and DINO trained on the HPA single-cell dataset (blue). E) Comparison of the HPA FOV protein localization classification models. X-axis: Number of parameters trained in a supervised fashion, with ground truth labels. Y-axis: Average F1-score in the HPA FOV protein localization competition. F) Same as in E, for the HPA single cell protein localization classification task.*

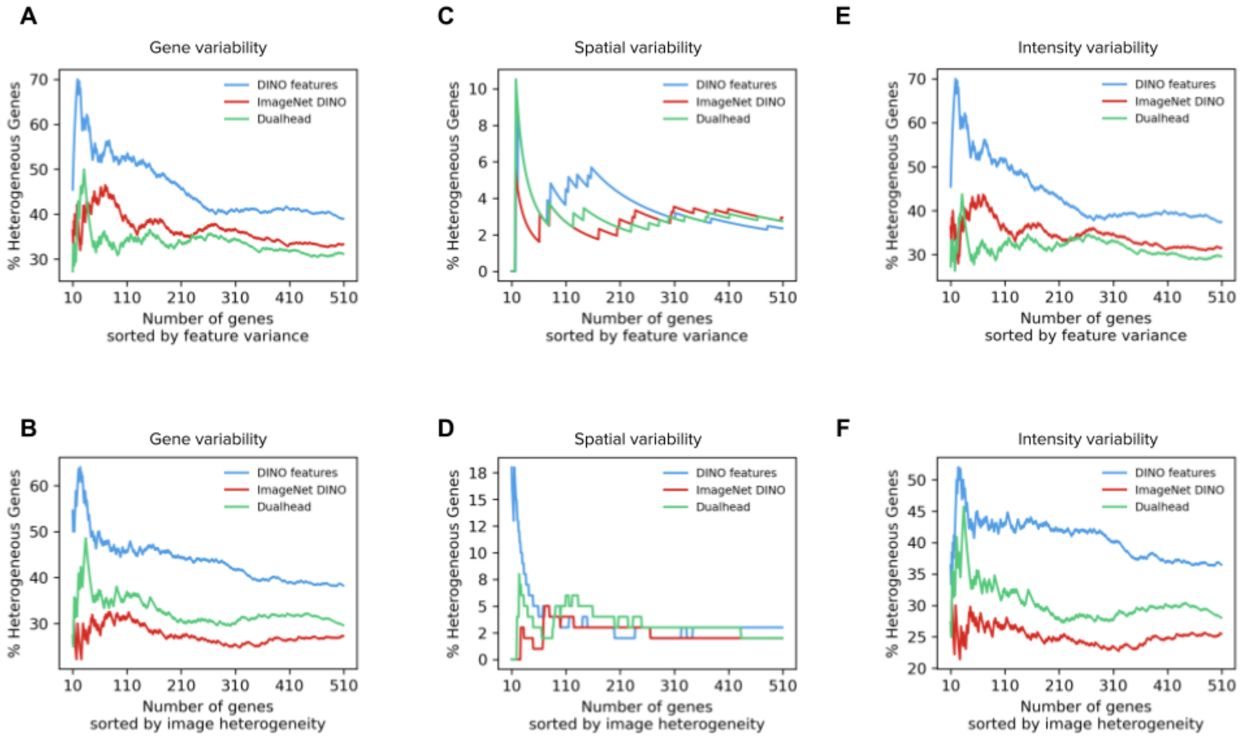


***Supplementary Figure S5. Protein localization heterogeneity in the HPA single-cell dataset, based on three models.*** *A) The x-axis represents the ranking of genes according to the variance of single-cell features in an image (Methods). The y-axis represents the proportion of genes labeled as heterogeneous according to existing annotations in the HPA website. The comparison is between three models: DINO trained on the HPA single cell dataset, DINO pretrained on ImageNet, and the Dualhead Kaggle group. B) Same as in A, with the genes ranked according to the standard variation of the single cell features across all images stained for the corresponding protein. C and D) Same as in A and B, with the ground truth being genes that were annotated for spatial variability. E and F) Same as in A and B, with the ground truth being genes that were annotated for intensity variability.*

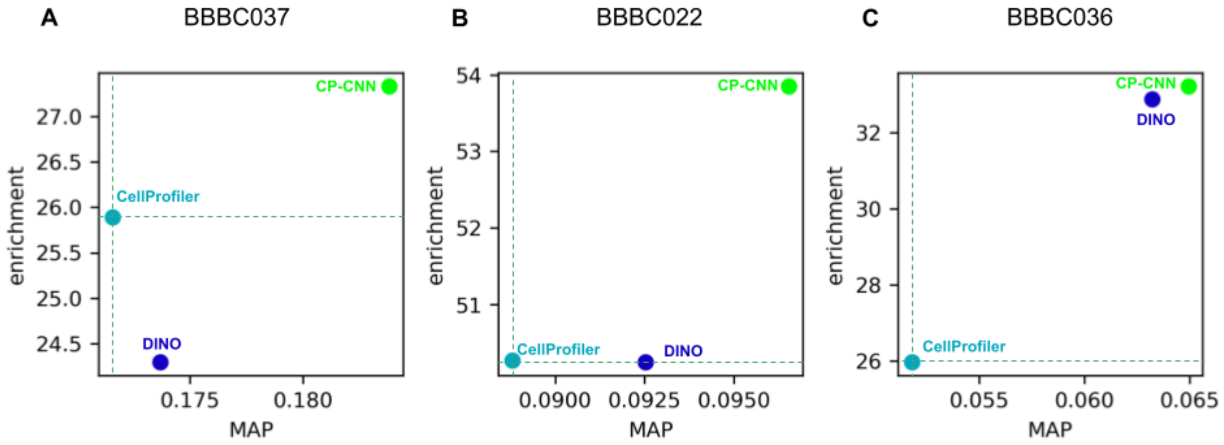


***Supplementary Figure S6.*** *A) Scatter plot comparing the performance of feature representations on the BBBC037 dataset. The x-axis shows the mean average precision of nearest-neighbor search queries to match mechanism-of-action classes. The y-axis shows the enrichment of the genes based on the ranking of the different features. B,C) Same as in B for the BBBC022 dataset, and the BBBC036 dataset, respectively.*
